# Multimodal lesion network mapping to predict sensorimotor behavior in stroke patients

**DOI:** 10.1101/2021.12.23.473973

**Authors:** Antonio Jimenez-Marin, Nele De Bruyn, Jolien Gooijers, Alberto Llera, Sarah Meyer, Kaat Alaerts, Geert Verheyden, Stephan P. Swinnen, Jesus M. Cortes

**Affiliations:** Computational Neuroimaging Group, Biocruces-Bizkaia Health Research Institute, Barakaldo, Spain.; Biomedical Research Doctorate Program, University of the Basque Country (UPV/EHU), Leioa, Spain.; Department of Rehabilitation Sciences, KU Leuven, Leuven, Belgium.; Movement Control and Neuroplasticity Research Group, Department of Movement Sciences, KU Leuven, Leuven, Belgium.; LBI-KU Leuven Brain Institute, Leuven, Belgium.; Department of Cognitive Neuroscience, Radboud University Medical Centre, Nijmegen, The Netherlands.; Centre for Cognitive Neuroimaging, Donders Institute for Brain, Cognition and Behavior, Nijmegen, The Netherlands.; Cell Biology and Histology Department, University of the Basque Country (UPV/EHU), Leioa, Spain.; IKERBASQUE, The Basque Foundation for Science, Bilbao, Spain.

**Keywords:** Lesion Network Mapping, Stroke, Canonical Correlation Analysis, Multimodal Imaging, Functional Magnetic Resonance Imaging, Diffusion Weighted Imaging.

## Abstract

Lesion network mapping (LNM) has proved to be a successful technique to map symptoms to brain networks after acquired brain injury. Beyond the characteristics of a lesion, such as its etiology, size or location, LNM has shown that common symptoms in patients after injury may reflect the effects of their lesions on the same circuits, thereby linking symptoms to specific brain networks. Here, we extend LNM to its multimodal form, using a combination of functional and structural connectivity maps drawn from data from 1000 healthy participants in the Human Connectome Project. We applied the multimodal LNM to a cohort of 54 stroke patients with the aim of predicting sensorimotor behavior, as assessed through a combination of motor and sensory tests. Test scores were predicted using a Canonical Correlation Analysis with multimodal brain maps as independent variables, and cross-validation strategies were employed to overcome overfitting. The results obtained led us to draw three conclusions. First, the multimodal analysis reveals how functional connectivity maps contribute more than structural connectivity maps in the optimal prediction of sensorimotor behavior. Second, the maximal association solution between the behavioral outcome and multimodal lesion connectivity maps suggests an equal contribution of sensory and motor coefficients, in contrast to the unimodal analyses where the sensory contribution dominates in both structural and functional maps. Finally, when looking at each modality individually, the performance of the structural connectivity maps strongly depends on whether sensorimotor performance was corrected for lesion size, thereby eliminating the effect of larger lesions that produce more severe sensorimotor dysfunction. By contrast, the maps of functional connectivity performed similarly irrespective of any correction for lesion size. Overall, these results support the extension of LNM to its multimodal form, highlighting the synergistic and additive nature of different types of imaging modalities, and the influence of their corresponding brain networks on behavioral performance after acquired brain injury.

## Introduction

Mapping the behavioral impact of brain lesions is critically important in clinical practice. Regardless of the cause of the lesion, its size or precise location, evidence has accumulated over recent years in favor of the connectome hypothesis whereby the specific network(s) the lesion affects can predict many of the patient’s responses across behavioral, cognitive or sensory-motor domains^1–22^. A computational solution to map the lesion-driven disconnectivity to behavior was developed recently^1^, and it has been successfully applied to several conditions and pathologies^4, 5, 7–13, 15, 16, 18, 19, 21–23^. Due to the simplicity of this method to correlate behavioral outcomes with the extent of lesion-driven disconnection, the strategy was referred to as Lesion Network Mapping (LNM). Here, we extend such LNM in to two strategic dimensions. Firstly, by proposing a multimodal approach in which we introduce a combination of both structural and functional networks to predict behavior. Secondly, and motivated by previous work^2, 6, 20^, by assessing behavioral performance using a combination of several multidomain scores.

We applied our novel strategy to stroke, a highly disabling condition triggered by a focal lesion that typically produces multiple behavioral deficits. Although stroke produces focal damage, it is well known to affect remote areas, such as regions directly connected to the lesion through long-range white matter tracts^24^ or indirectly connected regions. In terms of the latter, functional networks introduce indirect correlations through so-called common-neighbor interactions^25–27^. Significantly, the mapping of different behavioral deficits to imaging alterations localize tightly within specific brain networks^2, 6, 28, 29^. Indeed, the degree of network disruption was shown to be a good correlate of behavioral recovery from damage after stroke^30–32^, as also witnessed in longitudinal data^33–35^.

As stroke is a highly disabling condition, we assessed different aspects of the individual’s motor ability using two tests widely recognized as measures of motor performance: the Action Research Arm Test (ARAT) and the Fugl-Meyer assessment - upper extremity (FMA-UE) test^36, 37^. In terms of everyday activities that involve different movement modalities like grasping, grip and pinchforce, as well as gross movement, ARAT serves to measure upper limb dysfunction after stroke^36, 38, 39^. In addition, the FMA-UE test was used as a complementary assessment of motor dysfunction^40^. Somatosensory performance is also known to be highly impaired following stroke and it is generally associated with a deterioration in dexterity, manipulation abilities and bimanual hand coordination skills^41, 42^. To assess somatosensory capacity, we used the Erasmus-modified Nottingham Sensory Assessment (Em-NSA)^43, 44^ that evaluates tactile, proprioceptive and higher cortical somatosensation, along with the perceptual threshold of touch (PTT) test^45^ that principally assesses tactile function and that has been used previously in neuroimaging studies^46^.

Our main hypothesis here was that by uniquely extending LNM to its multimodal form, in which we include both functional (FC) and structural connectivity (SC) measures, it would facilitate a better understanding of the synergistic contributions of individual modalities to explain multi-domain behavioral outcomes in stroke patients. We also hypothesized that by applying the multimodal form of LMN, the variance explained in the brain maps would be enhanced by achieving greater coverage of the variation in multi-domain behavior. For this purpose, we employed a canonical correlation analysis (CCA) to link multi-domain behavior to different lesion connectivity maps.

## Methods

### Participants

In this study we included two cohorts of stroke patients previously assessed elsewhere^47, 48^. The first cohort consisted of 25 patients who developed upper limb sensorimotor impairments after stroke, and they were recruited from the University Hospital Leuven and the University Hospital St-Luc Brussels. The second cohort consisted of 29 stroke patients recruited at four different centers: UZ Leuven (Pellenberg), Jessa hospitals (Herk-de-Stad), Heilig Hart Hospital (Leuven) and RevArte (Antwerp). The pooling of these two cohorts enhanced the statistical power of the analyses. Thus, in total we analyzed 54 first-stroke patients (25 males) with a mean age of 68.78 (σ = 13.98, range 28 – 92 years), and a mean time interval between stroke and behavioral assessment of 25.61 days (σ = 20.32, range 4 – 64). The lesions were distributed evenly across the left and right hemispheres (27 in each) and the size of the lesions in the patients varied from 0.30 to 255.94 cm^3^. Due to the large variation in time between the stroke and behavioral assessment, and in the participant’s age and lesion size, these three variables were regressed out for further analyses. The behavior of both cohorts of patients was assessed at the hospital using a dedicated procedure not included in the daily clinical routine. Full details of the demographic and clinical characteristics of the participants in this study are given in Table 1. The study was approved by the Ethical Committees at the different Institutions from which the patients were recruited (codes S60278 and S54601) and all the participants provided their signed informed consent before enrolling on the study.

**Table 1.**
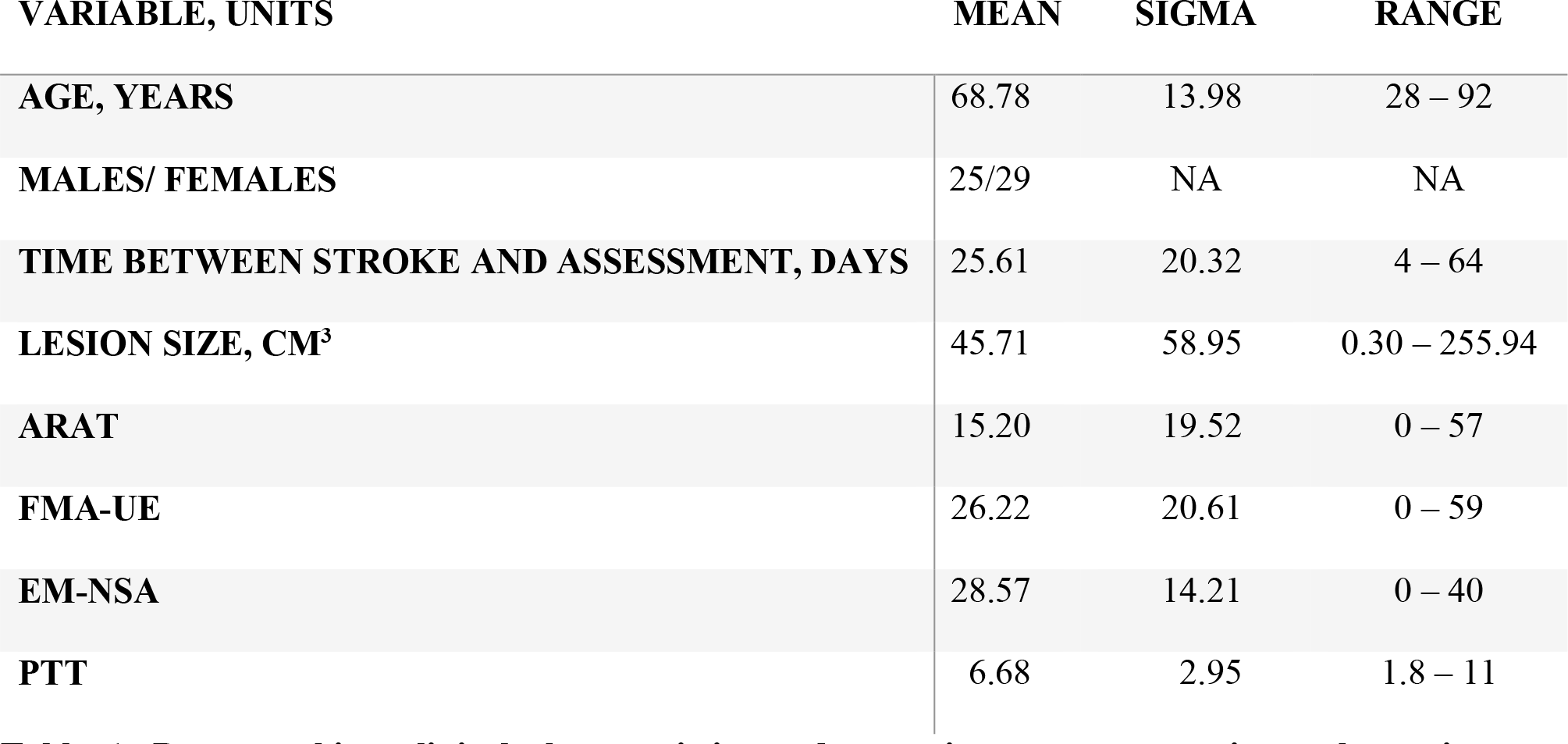
Demographics, clinical characteristics and sensorimotor outcomes in stroke patients. Abbreviations: ARAT - Action Research Arm Test; FMA-UE - Fugl-Meyer assessment for the upper extremity; EM-NSA - Erasmus modified Nottingham sensory assessment; PTT – perceptual threshold of touch.

| VARIABLE, UNITS | MEAN | SIGMA | RANGE |
| --- | --- | --- | --- |
| AGE, YEARS | 68.78 | 13.98 | 28 – 92 |
| MALES/ FEMALES | 25/29 | NA | NA |
| TIME BETWEEN STROKE AND ASSESSMENT, DAYS | 25.61 | 20.32 | 4 – 64 |
| LESION SIZE, CM<sup>3</sup> | 45.71 | 58.95 | 0.30 – 255.94 |
| ARAT | 15.20 | 19.52 | 0 – 57 |
| FMA-UE | 26.22 | 20.61 | 0 – 59 |
| EM-NSA | 28.57 | 14.21 | 0 – 40 |
| PTT | 6.68 | 2.95 | 1.8 – 11 |

### Image acquisition

T1 anatomical MRIs were acquired from all the stroke patients (N = 54) using a Philips 3T Achieva scanner equipped with a 32-channel head coil and applying the following parameters: 182 coronal slices covering the whole brain, repetition time (TR) = 9.6 ms, echo time (TE) = 4.6 ms, field of view (FOV) = 250×250 mm^2^, slice thickness = 1.2 mm and no interslice gap. FLAIR images were acquired with the following parameters: 321 transverse slices covering the whole brain, TR = 4800 ms, TE = 351 ms, inversion time = 1650 ms, FOV = 250 × 250 mm^2^, slice thickness = 1.12 mm and interslice gap = 0.56 mm. For the first cohort, we only acquired the FLAIR sequence. These images were used only for lesion segmentation.

We also analyzed images from healthy subjects (N=1000, ages ranging from 22 to 35 years old) obtained from the Human Connectome Project (HCP, WU-Minn Consortium, Principal Investigators David Van Essen and Kamil Ugurbil: 1U54MH091657) funded by the 16 NIH Institutes and Centers that support the NIH Blueprint for Neuroscience Research, and by the McDonnell Center for Systems Neuroscience at Washington University. For each HCP subject, MRI acquisition was performed using a 3T Siemens Connectome Skyra with a 100 mT/m and 32-channel receive coils. The acquisitions used for network disconnectivity analyses were: 1) A high-resolution anatomical T1-weighted 3D MPRAGE sequence with the parameters TR = 2400 ms, TE = 2.14 ms, Flip angle = 8 deg, FOV = 224×224 mm^2^, Voxel size = 0.7 mm isotropic, Acquisition time = 7 min and 40 s; 2) Functional data at rest to obtain the blood-oxygenation-level-dependent (BOLD) signals using a gradient-echo EPI sequence with the parameters TR = 720 ms, TE = 33.1 ms, Flip angle = 52 deg, FOV = 208 × 180 mm^2^, Matrix = 104 × 90, 72 slices per volume, a total number of 1200 volumes, Voxel size = 2 mm isotropic, Acquisition time = 14 min and 33 s; 3) Diffusion weighted data with a Spin-echo EPI sequence and the parameters TR = 5520 ms, TE = 89.5 ms, Flip angle = 78 deg, FOV = 210 × 180 mm^2^, Matrix = 168 × 144, 111 slices per volume, Voxel size = 1.25 mm isotropic, 90 diffusion weighting directions and six unweighted (b = 0) acquisitions, three shells of b = 1000, 2000 and 3000 s/mm^2^, Acquisition time 9 min 50 s. For further details on the acquisition parameters of the HCP participants see the documentation available at https://www.humanconnectome.org/.

### Image processing

#### Lesion segmentation

Lesion segmentation was based on both T1 and FLAIR images, and it was performed semi-automatically using the *clusterize* toolbox^49^ implemented in SPM12 and running in MATLAB R2019b, followed by manual inspection and correction in MRIcron by experienced researchers. After segmentation, the lesion masks were non-linearly co-registered to the MNI152 template with the dimensions 2×2×2 mm^3^. To enhance this co-registration we used the T1 sequence (or FLAIR in the first cohort), filling the lesioned area with healthy tissue from the contralateral hemisphere.

#### Functional images from healthy HCP participants

Resting state functional MRIs from HCP healthy controls (N=1000) were used to generate functional connectivity maps of lesions. First, the images were corrected for EPI gradient distortions and normalized to the MNI152 standard template with a voxel size equal to 2×2×2 mm^3^ using the HCP *fMRIVolume* and *fMRISurface* pipelines. After image normalization, we removed any nuisances with a procedure that mixes a volume-censoring strategy and a movement-related time-course regression, together with physiological signal regression. To do so, volumes were marked as censored when the frame-wise displacements (FDs) were greater than 0.2 or the derivative of the root-mean-squared variance was greater than 7.5%, following previous recommendations^50–52^. Moreover, the volume prior to, and the two following the censored one, were also marked as censored. The entire time series was then split into segments of 5 volumes in length, to finally remove all segments containing at least one contaminated volume, as well as the first segment. The 1000 subjects selected were those with the least contaminated volumes. Subsequently, any nuisances were removed while simultaneously applying a bandpass filter between 0.01-0.08Hz. Nuisance signals were the first five principal components of the CSF and white matter signals, the linear and quadratic trends, and the 24-parameter movement-related time-series. Finally, each filtered image was spatially smoothed with a Gaussian kernel of 6 mm FWHM.

After image preprocessing and to speed up computation, functional disconnection maps were obtained using only the first six minutes of the preprocessed 4D image. To do so, the lesion mask for each stroke patient was used as the seed for the analysis of seed-based connectivity (SBC), applied separately to each HCP subject. In such a way, the time-series of the BOLD signals were obtained from HCP data, while the stroke patients provided the seed for SBC. The Pearson correlation values, ‘𝑟‘, between the seed-time series (obtained by averaging all the voxel-time series within a given lesion) and all other voxel-time series in the brain, were Fisher-transformed by applying the inverse hyperbolic tangent of 𝑟, 𝑧 = 𝑎𝑟𝑡𝑎𝑛ℎ(𝑟). Therefore, for each HCP subject and stroke patient we obtained a 3D brain map of z-values. The final functional disconnection map per patient was obtained after one-sample T-test statistics were applied to the 1000 HCP different maps.

#### Diffusion weighted images from healthy HCP participants

Inspired by the strategy to obtain functional disconnection maps, we applied SBC to the diffusion data in order to obtain structural disconnection maps. We first made use of the *bedpostx*^53^ results obtained after applying the HCP pipeline to each subject. The Camino software (http://camino.cs.ucl.ac.uk/) was then used to obtain a deterministic tractography, with fiber assignment using a continuous tracking algorithm^54^ and all voxels within a given lesion as seeds for whole-brain fiber generation, employing a maximum curvature of 60° and a fractional anisotropy threshold of 0.15. The voxel-level fiber-counting maps were then binarized for each HCP subject, defining the extent to which a given voxel in the brain is connected to any voxel within the lesion. Finally, we obtained the final structural disconnection map by averaging across all HCP participants, one for each of the stroke patients in our cohort. Therefore, a given voxel in the final map had a value of 1 when the stroke patient’s lesion was connected to that voxel in all HCP participants and conversely, 0 if it was not connected in any HCP participant.

### Sensorimotor assessment as a behavioral outcome in stroke patients

Somatosensory performance was evaluated using the Em-NSA^43^ to assess exteroception, proprioception and higher cortical functions. The Em-NSA evaluates five distinct somatosensory modalities^55^ including light touch, pressure, pinprick, sharp-blunt discrimination and proprioception. Light touch was tested with cotton wool, pressure with an index finger pinprick with a toothpick, and sharp-blunt discrimination by alternating a toothpick prick of the index finger with that at the following contact points: fingers, hand, forearm, and upper arm. Proprioception was assessed during passive movements of the different upper limb joints. Each point of contact was assessed 3 times and graded on an ordinal scale as: 0, patient fails to detect any sensation on all 3 occasions; 1, patient identifies test sensation, but not on all 3 occasions; or 2, patient correctly identifies the test sensation on all 3 occasions. In total, the scores for each modality ranged from 0 (complete somatosensory impairment) to 8 (no somatosensory impairment). The total score for the Em-NSA (including all modalities together) ranges from 0 to 40, with a higher score representing better upper-limb somatosensory performance and a score below 36 indicating a degree of somatosensory impairment^38^. A second test that was used for somatosensory performance was the PTT^45^, which assesses gentle touch perception by applying transcutaneous electrical nerve stimulation (TENS) with a CEFAR Primo Pro apparatus (Cefar medical AB, Sweden). The scores reflect the mA applied to detect stimulation on the affected tip of the index finger and thus, higher PTT scores indicate more somatosensory impairment, with maximal stimulation was set at 10 mA to prevent burning. If a patient was unable to feel any sensation at the maximum stimulation level they were awarded a score of 11.

Motor performance was assessed with the ARAT to evaluate upper limb activity^39^, a 19-item test divided into 4 categories: grasp, grip, pinch, and gross arm movement. Each category is rated with an integer value ranging from 3 (patient performs test with normal motor pattern) to 0 (patient cannot perform any part of the test). Therefore, the maximum score after the ARAT test is 57, indicating normal motor performance of the arm. Motor performance was also assessed with the FMA-UE^40^ test, addressing motor function of the upper extremity as a whole (including shoulder, elbow, wrist and hand movements), from reflex activity to voluntary activation^55^. The total FMA-UE score ranges between 0 and 66, with a higher score representing better upper-limb motor function.

### Association between behavioral outcomes and the disconnectivity maps

#### Data pre-processing

Because there were some missing values in the behavioral scores from stroke patients, we applied the following procedure to this data. Rather than penalizing the sample size by eliminating these patients, we generated the missing values using an iterative imputer algorithm with extra tree regressors, seen to be highly effective in generating missing data^56^. The total number of missing values was zero for the ARAT test, 3 for the FMA-UE test, 2 for the Em-NSA and 2 for the PTT.

For the disconnectivity maps, we generated a binary mask for each modality, attributing a value of ‘1’ for the voxels that exhibited disconnection (or an existing lesion in the case of the lesion modality), and ‘0’ otherwise. We next built a matrix for each modality of the dimension (# of stroke patients) x (# of voxels) in each modality mask. Finally, to reduce the dimension of this matrix, we employed a principal component analysis (PCA), keeping a different number of components for each mask to maximize performance and using it for subsequent imaging-behavior association analysis.

#### Canonical correlation analysis (CCA)

To establish associations between the imaging and behavioral outcomes, we applied an iterative CCA. For this, the residuals of the behavioral variables were used as the dependent variables (after regressing out lesion size, patient age, and time between stroke and assessment) and the PCA components of the disconnectivity maps as the independent variables, starting with one component as the X variable and ending with a matrix including all the principal components. When the number of components in X increased, the CCA strategy led to overfitting, making the extent of the correlation achieved meaningless. To overcome this limitation, we performed a predictive CCA approach using leave-one-out cross-validation (LOOCV). Thus, to obtain the CCA correlation value, for every step (subject) we adjusted the CCA with *all-except-one* participants and calculated the canonical scores of the remaining participant with the *learned model*. After completing the loop over subjects we obtained ‘predicted’ canonical scores for all the subjects and a ‘predictive’ CCA score was computed. Statistical significance was assessed by surrogate generation of 1000 random permutations of the X variables and the p-value calculated as the number of instances where the surrogate correlations were greater than those produced by CCA divided by the total number of permutations (i.e.: permutation testing)^57^. Finally, we selected the *best* model as that for which the PCA order achieved the maximum correlation *T* using the predictive LOOCV-CCA introduced.

This procedure was performed for SC and FC separately (unimodal analyses), and combined through spatial concatenation of the SC and FC matrices (multimodal analysis). This analysis was followed by a PCA and the *mixed* components obtained were used to apply a CCA, as for the unimodal cases. In the multimodal approach, we first transformed the SC to logarithmic values, and then standardized both the SC and FC matrices before concatenation.

#### Brain maps corresponding to *best* CCA solutions

Using the weights of the *best* model and their coefficients, we projected this solution back onto the brain space. The final maps were obtained by transforming the values to z-scores and representing only the 𝑍 > 2 values. For the multimodal strategy, we first back-projected the best solution, and then we split the different coefficients identifying the separate SC and FC contributions.

#### Neurobiological description of the brain maps

Brain maps were described using a combination of four different atlases. The first one was the XTRACT atlas^58^ that is composed of 42 different white matter tracts, including: 10 association tracts (L/R hemispheres), 4 commissural tracts, 4 limbic tracts (L/R hemispheres), and 5 projection tracts (L/R hemispheres). The second was the Desikan-Killiany partition^59^ that has 88 regions, 8 of them subcortical (L/R hemisphere) and 36 cortical (L/R hemispheres). To this atlas, we added the brainstem as an additional region. The third atlas was the SMATT atlas^60^ with 60 regions in total, 30 sensorimotor tracts (L/R hemispheres). Finally, the fourth atlas was an overlay of several partitions proposed by Yeo and collaborators, and that included the cortex^61^, cerebellum^62^, striatum^63^ and thalamus (to the best of our knowledge released by the author but not yet published in any citable reference).

### Statistical analyses

Unless otherwise specified, group comparisons of different metrics used in this study (e.g., scores in patients with a lesion in the right hemisphere compared to those in the left hemisphere) were performed with two-sample Kruskal-Wallis non-parametric tests. Statistical dependencies between behavioral scores were assessed through a Pearson correlation analysis. Spatial similarity between brain maps was assessed using a Pearson spatial correlation.

## Results

Multimodal LNM was applied to a cohort of first-time stroke patients with sensory-motor impairments (N = 54). Behavior alterations were evaluated with a battery of sensorimotor tests, the demographic, clinical and sensorimotor scores and details of which are seen in Table 1.

After projecting all the patient’s lesions onto the same template (MNI152 2 mm³), we obtained the functional and structural disconnection maps for each patient following the pipeline detailed in Figure 1. This pipeline also made use of functional and structural imaging data from healthy participants from the HCP (N=1000). In other words, our method analyzed the impact that the patient’s lesion had on the disconnection of specific networks that exist in healthy brains. At the population level (N=54 stroke patients), we concatenated the different patient’s disconnection maps into a final matrix and applied a PCA to get the principal components that were then used as independent variables in a CCA.

**Figure 1.**
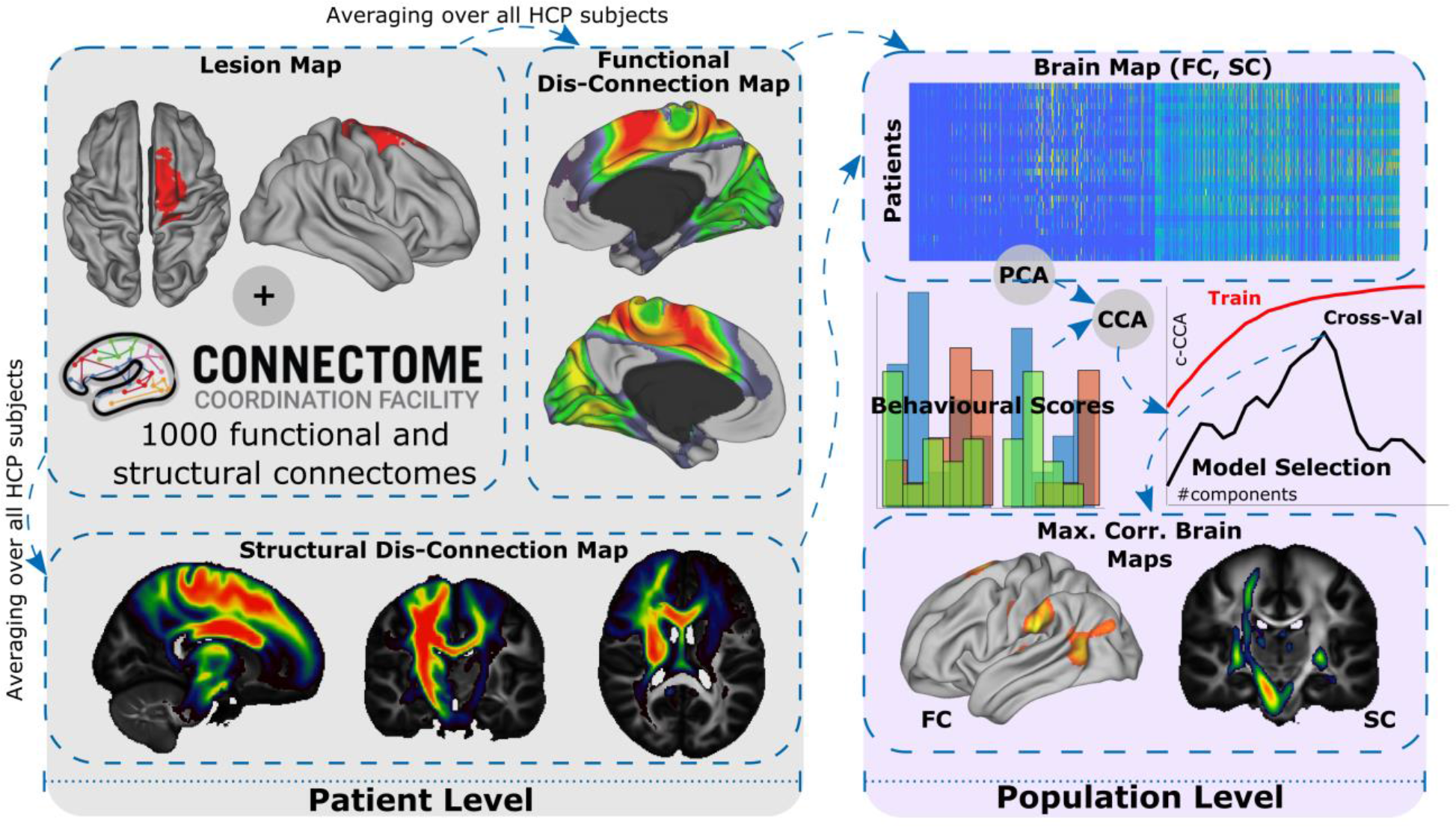
Pipeline for multimodal lesion network mapping and its association to behavioral outcome after stroke through Canonical Correlation Analysis (CCA). At the patient level (gray shading), brain lesion masks are used as seed regions to calculate the functional correlation maps (applying seed-based correlation analysis and using the segmented lesion as the seed for each HCP subject) and the structural correlation maps (applying tractography from the segmented lesion to the rest of the brain for each HCP subject) from a group of healthy control participants from HCP (N=1000). After averaging all the participants in the HCP dataset (see Methods for details), we obtained the functional disconnection maps for each patient, accounting for the functional impact of lesion disconnection, and likewise for the structural disconnection maps. At the population level (purple shading), a matrix with dimensions (# of stroke patients) x (# of voxels) per modality map (FC or SC) was built and reduced using a PCA, which returns a new matrix with (# of patients) x (# of principal components) dimensions, the PCA components considered here as the brain map features. The association between the features of the SC and FC, and the behavioral scores was obtained by applying a CCA. As the number of features increases, the correlation between features and behavior (represented here as c-CCA) increases up to values close to 1 (red curve, Train), dealing with overfitting. Cross-validation techniques can overcome this problem (for details see Methods). For the maximum CCA correlation value in the cross-val curve (black), we built brain maps of those components producing maximum performance. The maps can be obtained in a single modality, here shown for FC or SC, or as a combination of them (not shown here but implemented in this study). Abbreviations: FC, functional connectivity; SC, structural connectivity; PCA, principal component analysis; CCA, canonical correlation analysis.

The dependent variables for the CCA were the behavioral sensorimotor scores. The two ARAT and FMA-UE motor scores were correlated to each other (*r* = 0.85, *p* < 0.001: Figure 2), as were the sensory Em-NSA and PTT variables (*r* = -0.7, *p* < 0.001). However, the correlations between motor and sensory variables were not significant^†^, suggesting linear independence between the two motor and somatosensory domains.

**Figure 2.**
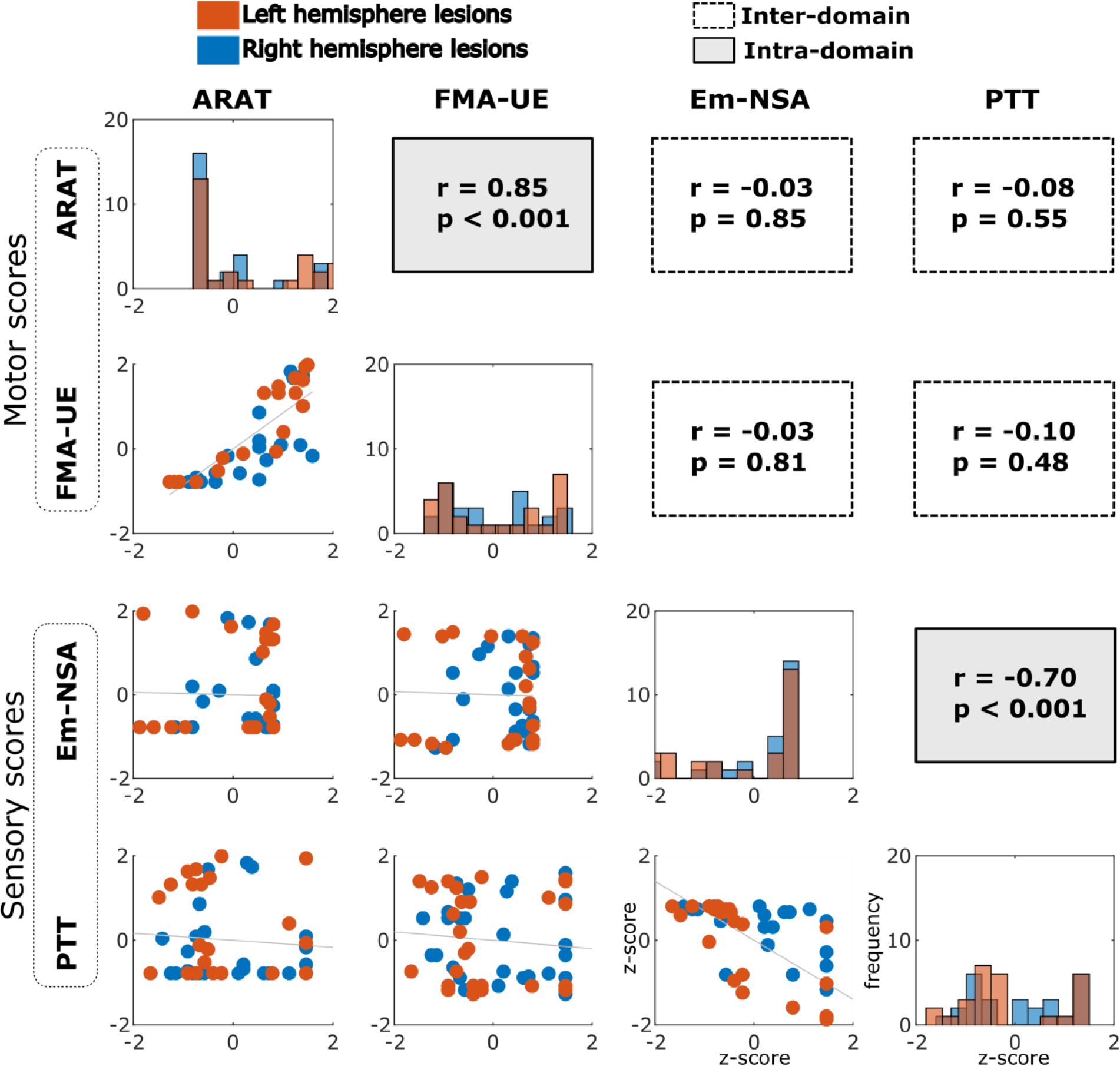
Distribution of behavioral -- motor and sensory -- scores. The principal diagonal panels represent the histogram values for each of the four behavioral scores, two being motor scores (ARAT and FMA-UE) and two somatosensory scores (Em-NSA and PTT). Off-diagonal panels (below the diagonal) show scatter plots between pairs of scores. We also provided Pearson correlation values (r) and associated p-values above the diagonal in the off-diagonal panels. The Red and blue colors represent scores from patients having lesions in the left and right hemispheres, respectively. Because the two datasets distribute equally well for the different scores, the left and right lesion datasets were pooled into a single cohort in this study. All the scores are represented here as Z-scores. Abbreviations: ARAT, Action Research Arm Test; FMA-UE, Fugl-Meyer Assessment; Em-NSA, Erasmus modified-Nottingham Sensory Assessment; PTT, Perceptual Threshold of Touch.

In addition, the differences in the scores between patients whose lesion was found on the left hemisphere as opposed to those who had it in the right hemisphere were not statistically significant^‡^. Indeed, there was strong similarity between the lesion brain maps in the left hemisphere and those of the right (*r* = 0.71, *p* < 0.001: Figure S1). Both these factors (the non-significant differences between behavioral scores in patients with a lesion in the left or right hemisphere, and the strong similarity between the two maps) justified merging the patients with lesions in the left or right hemisphere into a single cohort, thereby increasing the statistical power for our CCA analysis.

We next analyzed the statistical relationship between the behavioral scores and the confounders age, time between stroke and the behavioral assessment, and lesion size (Figure S2). The two latter variables showed significant correlations with the somatosensory outcome assessed by Em-NSA (*r* = 0.32, *p* = 0.02 and *r* = -0.45, *p* < 0.001, respectively), while lesion size was correlated with the PTT score (*r* = 0.42, *p* = 0.002).

We continued applying a CCA to assess the association between lesion connectivity maps and behavioral performance. After sequentially increasing the number of PCA components, the *best* model was chosen from the cross-validated curve (black curve in Figure S3), selecting the number of components (#Comp) that produce the maximum significant correlation (T). The resulting brain maps (𝑍 > 2) are illustrated in Figure 3 for both the unimodal (FC and SC) and multimodal analyses (FC + SC). Moreover, the 𝑍_𝑚𝑎𝑥_ value and the amount of variance explained (Ve) in #Comp were also evident for each case (Figure 3).

**Figure 3.**
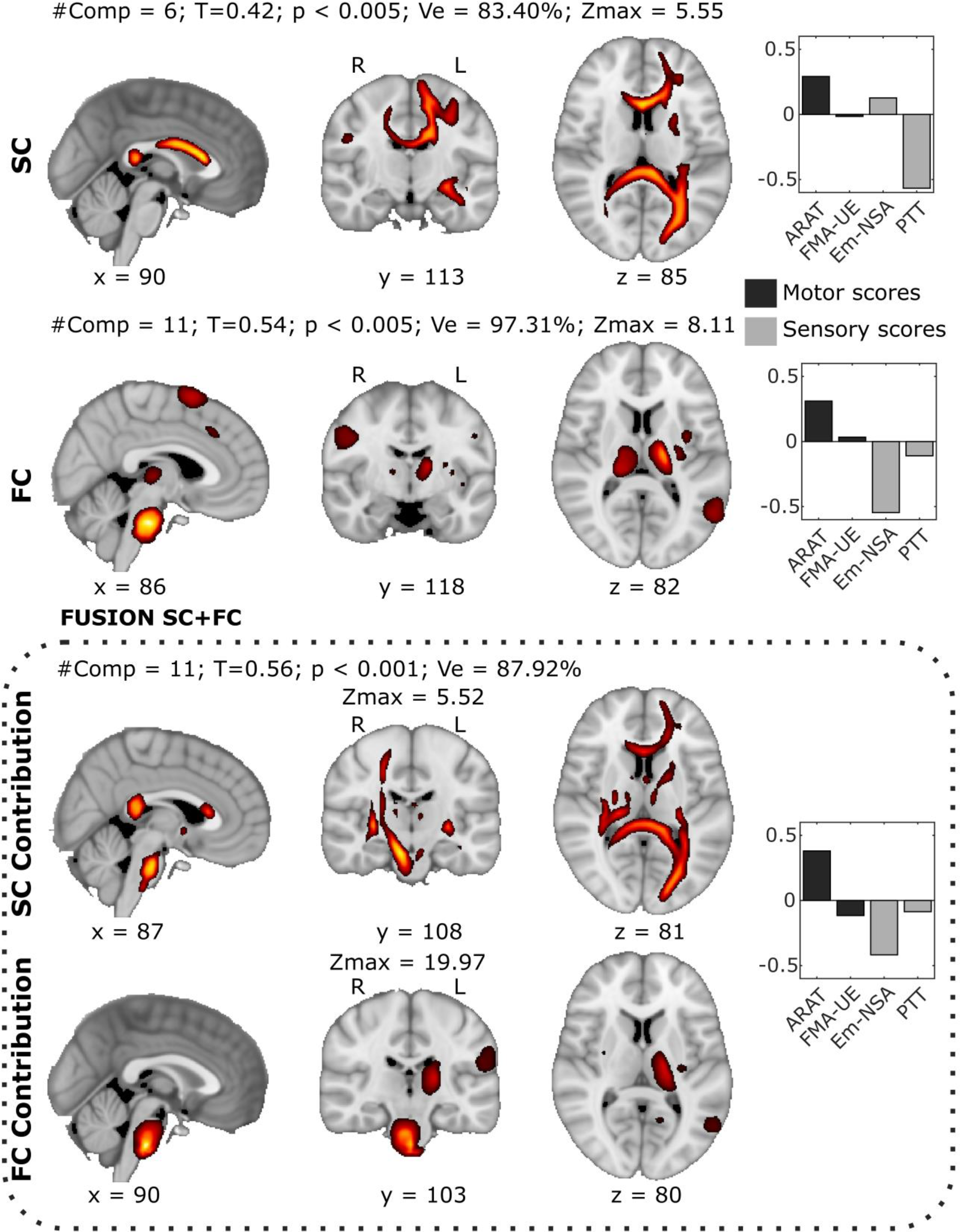
Brain maps with a maximal behavioral association from the unimodal and multimodal CCA after regressing out the lesion size effect. Final maps of the CCA solution that provide the maximum correlation between the X variables (the PCA components from each modality) and the Y variables (a combination of several behavioral scores represented in Figure 2). From top to bottom, SC brain maps (accounting for SC disconnectivity), FC brain maps, and SC+FC brain maps (box with dashed line). Moreover, the bottom panel also shows the individual SC and FC contributions to the maximum performance achieved by the multimodal SC+FC strategy. Together with the maximum behavioral-association maps, in all cases we provide the number of PCA components used (#Comp), the maximum correlation value (T), p-value (p), the amount of variance explained (Ve) and the maximum Z-value depicted in each map (Zmax). For visualization, all maps were threshold to 𝑍 > 2). In the left panel of each row we represent the behavioral weights corresponding to the maximum behavioral-association solution. Abbreviations: ARAT, Action Research Arm Test; FMA-UE, Fugl-Meyer Assessment; Em-NSA, Erasmus modified-Nottingham Sensory Assessment; PTT, Perceptual Threshold of Touch; FC, functional connectivity; SC, structural connectivity.

From the SC brain maps (see Tables S1-S2), the major tracts participating in the two unimodal and multimodal analyses were seen to be the forceps major, left frontal aslant tract, left anterior thalamic radiation, bilateral superior longitudinal fasciculus and bilateral optic radiation. Importantly, some tracts only in the brain only appeared to participate after the multimodal analyses, specifically, the right corticospinal tract and the middle cerebellar peduncle. Next, when looking at the overlap between SC maps and major sensorimotor-tracts represented in the SMATT atlas (a more specific sensorimotor atlas, cf. Tables S3-S4), unimodal SC maps included the left dorsal and ventral premotor cortices, right pre-supplementary motor and right primary somatosensory cortex. By contrast, multimodal SC maps included the right primary motor area, right primary somatosensory cortex and left ventral premotor cortex. When the significant association (Z > 2) between the two hemispheres was assessed (L and R), we found significant differences in the number of voxels between the unimodal analysis (L, N = 12203; R, N = 2589 and the multimodal approach (L, N = 7450; R, N = 5606). Hence, there was a significant shift from the left to right hemisphere when going from unimodal to multimodal maps (*χ*^2^ = 2160.04, *p* < 0.001).

When looking at the FC maps (Tables S5-S6), the brain regions that participated in both unimodal and multimodal associations were the brainstem, left supramarginal nucleus (the part of it that overlaps with the secondary somatosensory cortex), left thalamus, bilateral superior frontal cortex (overlapping with the premotor cortex), left inferior parietal and right precentral cortex (overlapping with the primary motor cortex and primary sensory cortex). More specifically, when looking at the overlap between the FC maps and the major Resting State Networks (RSNs: Tables S7-S8), the unimodal FC maps overlapped with the dorsal attention and limbic networks, while a major participation for ventral attention and sensorimotor networks was evident in the multimodal FC analysis. Similar to the SC, when the activation in the two hemispheres was compared we found N = 4023 (L) and N = 3269 (R) in the unimodal analysis, and N = 3369 (L) and N = 1958 (R) for the multimodal approach, confirming a significant shift from the right to the left hemisphere when going from unimodal to multimodal (*χ*^2^ = 82.70, *p* < 0.001), the opposite direction to the SC.

The behavioral weights corresponding to the *best* model solution were assessed (Figure 3). Strikingly, as opposed to the multimodal analyses our results for the unimodal analyses showed an imbalance in the optimal weights, with greater weights assigned to the somatosensory than the motor domain. By contrast, the multimodal solution provided a more balanced state between motor and somatosensory contributions. Moreover, the individual contributions of the SC and FC within the multimodal *best* solution had different strengths, as witnessed by the median value of the weight distribution for the SC (0.08) and FC (0.12), and indicating a major participation of the FC as opposed to the SC in the multimodal solution. Furthermore, when comparing the performance of the behavioral association between the unimodal and multimodal analyses, the FC outperformed SC at approximately 30% higher values of maximum correlation and at 16% higher values of explained variance.

Finally, in an attempt to address which modality was most sensitive to eliminate the effect of lesion size, we repeated a similar strategy based on lesion connectivity maps but without removing this effect (Figure S4). Notably, while the white matter tracts that participated in the maximum CCA solution were similar between the two approaches (see Figure 3 and Figure S4), the SC performance was nevertheless much better without removing the lesion size (no lesion size removal T = 0.63, lesion size removed T = 0.42). However, the removal of lesion size did not affect the FC performance. We performed a similar analysis but for lesion maps with and without the removal of lesion size (Figure S5). As with the SC but unlike the FC, the performance for lesion maps was also much better when the lesion size effect was removed (T = 0.61 vs. T = 0.38). These results indicate that both the SC disconnection maps and lesion maps are more sensitive to lesion characteristics than the FC maps.

## Discussion

Lesion network mapping (LNM) is a novel technique based on brain connectivity that has significant clinical impact, a conceptually straightforward approach that has been shown to perform impressively^64^. Critically, LNM makes use of human brain lesion models but it only takes into account the position of the lesion, reconstructing which brain regions in a healthy population of young adults (N = 1000) that are close to or at a distance from the lesion putatively become disconnected in the patient. In this way, LNM serves to stratify lesion-triggered symptoms through differentiated network mapping, offering two direct advantages over other *patient-specific* brain connectivity approaches. From a technical perspective, the reconstruction of networks in lesioned brains remains a challenge^65^, but this is not necessary for LNM. From a clinical point of view, LNM is not demanding as lesion segmentation can be performed after acquiring just a few FLAIR or T2 slices, as opposed to having to follow a complete multimodal MRI protocol, although for some conditions segmentation might be also challenging^66^.

Our contribution here further extends the use of LNM in two critical directions. First, by combining FC and SC networks we assess the multimodal impact of a lesion on each of these, through which the incorporation of other modalities would be straightforward, such as different PET classes. Second, we study the association between multimodal networks and multi-domain sensorimotor behavior, since the dysfunctionality caused by stroke cuts across different behavioral spheres. Therefore, a systemic characterization of patients requires the use of methods that can deal with this behavioral complexity. In the present work, these two novel multimodal and multi-domain aspects are combined using a CCA, a paradigm to map the images onto behavior, thereby leveraging the clinical relevance of these studies and their fundamental interest in neuroscience^67–70^.

Our multimodal LNM data highlights several improvements when compared to the unimodal approach. The multimodal analysis reveals that functional maps contribute more strongly than structural maps to the optimal prediction of sensorimotor behavior. This might reflect the fact that it is easier to adjust FC than SC after the insult, leaving room for larger FC variations to match the more extensive behavioral repertoire during recovery. Indeed, this is possible because FC changes occur over a more rapid time scale than those in SC. In relation to the number of behavioral domains covered by the best solution to link brain maps and behavior, we found that while both unimodal SC and FC better represent the sensory domain, the multimodal LNM provided a more balanced sensorimotor representation, supporting the notion that multidomain behavior could impose a major imprint on multimodal circuits. Our results suggest a possible mechanism to achieve this, as specifically reflected by the fact that in the multimodal analyses the FC brain maps shift from the right to the left hemisphere, whilst the SC shift in an opposite direction, from left to the right, thereby increasing the spatial coverage within the brain. This might be related to the contralateral lesion activity observed after stroke^71^. Finally, the multimodal analysis showed that the shared variance (across functional and structural data at the patient level) reveals the importance of regions and networks that did not appear in the unimodal analyses, such as the corticospinal tract (CST) and the primary motor area (M1), both structures well-known to be critical for motor function. In addition, the involvement of the middle cerebellar peduncle was evident, which mediates the communication between the cerebellum and the prefrontal cortex in the coordination and planning of motor tasks^72^. Moreover, the networks emerging from the multimodal analysis were the ventral attention network (VAN) and sensorimotor network, the former known to be relevant for stimulus driven attention, such as when somatosensory input is being processed^73^.

Another issue to highlight is what happened when we removed the effect of lesion size from our analyses. Unimodal SC and lesion maps were highly dependent on this manipulation and their performance deteriorated dramatically after lesion size was removed, as witnessed here by the amount of variance explained and the maximum correlation with behavior achieved. By contrast, FC provided similar performance irrespective of any correction for lesion size, suggesting that the FC is less localized (or more redundant) when processing sensorimotor tasks relative to the structural maps.

An interesting observation from our data was that lesion size was significantly correlated with the two sensory tests (Em-NSA and PTT) but not with either of the two motor tests (ARAT and FMA-UE). This suggests that the motor outcome assessed in this study was apparently processed in more localized brain regions that, once affected, would provoke a worse outcome regardless of the lesion size. Conversely, because lesion size was correlated with sensory outcomes, it suggests that the sensory consequences of the lesion are distributed more broadly across circuits encompassing different brain areas (visual – parietal – frontal), in agreement with previous results showing a stronger interaction between brain areas for sensory processing than for motor tasks^74^. This phenomenon might also be related to the amount of cognitive intervention required. While motor outcomes can be processed in a more straightforward manner, the sensory outcomes require the recruitment of more resources involved in cognitive control, including those required for attention deployment, although this possibility clearly requires further clarification.

Recent work acknowledged certain limitations to LNM^75, 76^. For example, when dealing with large lesions containing both white and grey matter, using the segmented lesion as the only seed to perform LNM can introduce relevant methodological biases due to the differences in the BOLD signal between voxels belonging to gray or white matter. To address FC, here we adopted a more fine-grained approach by averaging the signals within the gray matter. Moreover, the stroke patients studied were recruited paying attention to whether they had sensory impairment and independently whether they suffered any effect in motor performance, which might introduce some bias with respect to other studies in which patients with greater motor dysfunction were recruited.

In conclusion, by applying our multimodal LNM approach to predict multidomain sensorimotor behavior, we present evidence of the synergistic and additive influence of different types of brain networks on patients after brain injury, affecting their outcome and thereby making the whole more than the sum of its parts. Moreover, when a patient’s behavior is assessed across multiple domains of cognitive, sensory and motor function, our methodological approach, combining structural and functional maps, appears to be the most suitable when assessing such multidomain outcomes.

## Supplementary material

**Table S1.** Anatomical description of unimodal SC brain maps corresponding to the CCA solution and providing maximum association with sensorimotor behavior. Labels correspond to the XTRACT atlas, and only tracts attributed Z > 2 and with a percentage overlap > 50 voxels are shown.

| Tract | Peak x | Peak y | Peak z | Zmax | Nvoxels | Perc. Overlap |
| --- | --- | --- | --- | --- | --- | --- |
| *Forceps Major* | -8 | -34 | 12 | 4.66 | 1157 | 39.54 |
| *Frontal Aslant Tract L* | -16 | 8 | 30 | 4.76 | 890 | 60.26 |
| *Superior Thalamic Radiation L* | -16 | 8 | 44 | 4.41 | 875 | 44.30 |
| *Superior Longitudinal Fasciculus 1 L* | -16 | -4 | 42 | 4.67 | 731 | 31.56 |
| *Optic Radiation L* | -28 | -30 | 4 | 4.20 | 668 | 71.14 |
| *Superior Longitudinal Fasciculus 2 L* | -28 | 24 | 26 | 3.37 | 645 | 23.59 |
| *Inferior Fronto-Occipital Fasciculus L* | -26 | -52 | 20 | 4.63 | 482 | 30.51 |
| *Middle Longitudinal Fasciculus L* | -28 | -58 | 20 | 4.53 | 452 | 33.28 |
| *Anterior Thalamic Radiation L* | -18 | 32 | 22 | 3.87 | 414 | 22.66 |
| *Fornix L* | -10 | -34 | 12 | 4.50 | 411 | 40.69 |
| *Forceps Minor* | -12 | 22 | 16 | 5.48 | 358 | 19.47 |
| *Cingulum subsection: Dorsal L* | -10 | 0 | 30 | 4.60 | 314 | 22.49 |
| *Acoustic Radiation L* | -26 | -28 | 4 | 3.91 | 214 | 31.01 |
| *Frontal Aslant Tract R* | 16 | 14 | 26 | 4.26 | 212 | 13.55 |
| *Corticospinal Tract L* | -12 | -20 | 58 | 3.19 | 192 | 9.01 |
| *Superior Longitudinal Fasciculus 1 R* | 12 | 6 | 24 | 4.02 | 188 | 7.67 |
| *Superior Thalamic Radiation R* | 18 | -2 | 38 | 3.43 | 139 | 7.18 |
| *Cingulum subsection: Temporal L* | -24 | -42 | -2 | 2.82 | 137 | 16.47 |
| *Vertical Occipital Fasciculus L* | -18 | -82 | 18 | 2.96 | 126 | 8.36 |
| *Anterior Thalamic Radiation R* | 20 | 30 | 22 | 3.75 | 96 | 5.08 |
| *Uncinate Fasciculus L* | -36 | -6 | -20 | 2.96 | 93 | 8.32 |
| *Anterior Commissure* | -28 | -8 | -10 | 3.55 | 88 | 28.12 |
| *Arcuate Fasciculus L* | -32 | -34 | 18 | 2.80 | 75 | 5.34 |
| *Inferior Longitudinal Fasciculus L* | -40 | -16 | -20 | 2.89 | 73 | 3.47 |
| *Optic Radiation R* | 24 | -44 | 24 | 3.19 | 52 | 6.25 |

**Table S2.** Anatomical description of the multimodal SC brain maps corresponding to the CCA solution and providing maximum association with sensorimotor behavior. Labels correspond to the XTRACT atlas, and only tracts attribute Z > 2 and with a percentage overlap > 50 voxels are shown.

| Tract | Peak x | Peak y | Peak z | Zmax | Nvoxels | Perc. Overlap |
| --- | --- | --- | --- | --- | --- | --- |
| *Forceps Major* | -8 | -34 | 12 | 4.68 | 1720 | 58.78 |
| *Corticospinal Tract R* | 8 | -22 | -26 | 5.46 | 779 | 35.47 |
| *Anterior Thalamic Radiation L* | -6 | -16 | 8 | 3.69 | 560 | 30.65 |
| *Middle Cerebellar Peduncle* | 2 | -20 | -32 | 4.16 | 541 | 19.77 |
| *Frontal Aslant Tract L* | -14 | 22 | 38 | 3.03 | 485 | 32.84 |
| *Optic Radiation L* | -24 | -76 | 14 | 4.27 | 467 | 49.73 |
| *Forceps Minor* | -10 | 24 | 12 | 4.61 | 434 | 23.60 |
| *Optic Radiation R* | 24 | -50 | 30 | 4.39 | 357 | 42.91 |
| *Superior Longitudinal Fasciculus 1 L* | -22 | -56 | 34 | 3.11 | 345 | 14.90 |
| *Middle Longitudinal Fasciculus R* | 26 | -50 | 28 | 4.56 | 320 | 23.43 |
| *Middle Longitudinal Fasciculus L* | -24 | -74 | 16 | 4.27 | 264 | 19.44 |
| *Fornix L* | -10 | -34 | 12 | 4.55 | 253 | 25.05 |
| *Superior Longitudinal Fasciculus 1 R* | 24 | -58 | 32 | 3.87 | 242 | 9.88 |
| *Cingulum subsection: Dorsal L* | -12 | 28 | 16 | 3.98 | 216 | 15.47 |
| *Anterior Commissure* | 28 | -12 | -4 | 4.98 | 190 | 60.70 |
| *Inferior Fronto-Occipital Fasciculus R* | 30 | -18 | 2 | 5.34 | 181 | 10.36 |
| *Inferior Fronto-Occipital Fasciculus L* | -26 | -52 | 18 | 4.70 | 173 | 10.95 |
| *Vertical Occipital Fasciculus L* | -18 | -82 | 18 | 3.85 | 154 | 10.21 |
| *Superior Thalamic Radiation R* | 22 | -18 | 44 | 3.78 | 145 | 7.49 |
| *Acoustic Radiation R* | 32 | -28 | 8 | 4.12 | 108 | 16.69 |
| *Corticospinal Tract L* | -26 | -16 | -2 | 4.22 | 98 | 4.60 |
| *Superior Longitudinal Fasciculus 3 R* | 38 | -2 | 28 | 2.78 | 78 | 3.92 |
| *Acoustic Radiation L* | -28 | -24 | 4 | 4.82 | 77 | 11.16 |
| *Superior Longitudinal Fasciculus 2 R* | 36 | -4 | 32 | 2.89 | 65 | 2.51 |
| *Superior Longitudinal Fasciculus 2 L* | -30 | -44 | 28 | 2.33 | 63 | 2.30 |

**Table S3.** Percentage overlap between the unimodal SC brain maps and the major sensorimotor brain tracts. The labels correspond to the SMATT atlas, and only tracts attributed Z > 2 and with a percentage overlap > 10 voxels are shown.

| *Tract* | *Peak x* | *Peak y* | *Peak z* | *Zmax* | *Nvoxels* | *Perc. Overlap* |
| --- | --- | --- | --- | --- | --- | --- |
| *Left-PMd* | -14 | -6 | 56 | .88 | 201 | 72.30 |
| *Left-PMv* | -14 | 10 | 50 | .17 | 196 | 70.76 |
| *Right-M1, PMd, PMv, SMA, preSMA, S1* | -18 | -10 | 56 | .53 | 167 | 66.53 |
| *Right-M1, PMd, PMv, SMA, preSMA* | -30 | -18 | 62 | .01 | 116 | 19.66 |
| *Right-preSMA* | 18 | 0 | 42 | .18 | 52 | 18.51 |
| *Right-S1* | 44 | -24 | 48 | .48 | 52 | 17.93 |
| *Left-M1, SMA* | -16 | -4 | 58 | .53 | 52 | 61.91 |
| *Right-SMA* | 18 | -6 | 44 | .28 | 51 | 17.59 |
| *Left-PMd, SMA* | -14 | 0 | 58 | .63 | 39 | 79.59 |
| *Left-M1* | -22 | -4 | 30 | .64 | 25 | 7.31 |
| *Right-PMd* | 20 | -10 | 46 | .44 | 11 | 4.35 |
| *Right-SMA, preSMA* | 18 | -2 | 44 | .25 | 11 | 28.95 |
| *Left-PMd, PMv* | -20 | 2 | 44 | .17 | 11 | 64.71 |
| *Left-M1, PMd, SMA, S1* | -18 | -4 | 52 | .38 | 11 | 22.45 |

**Table S4.**
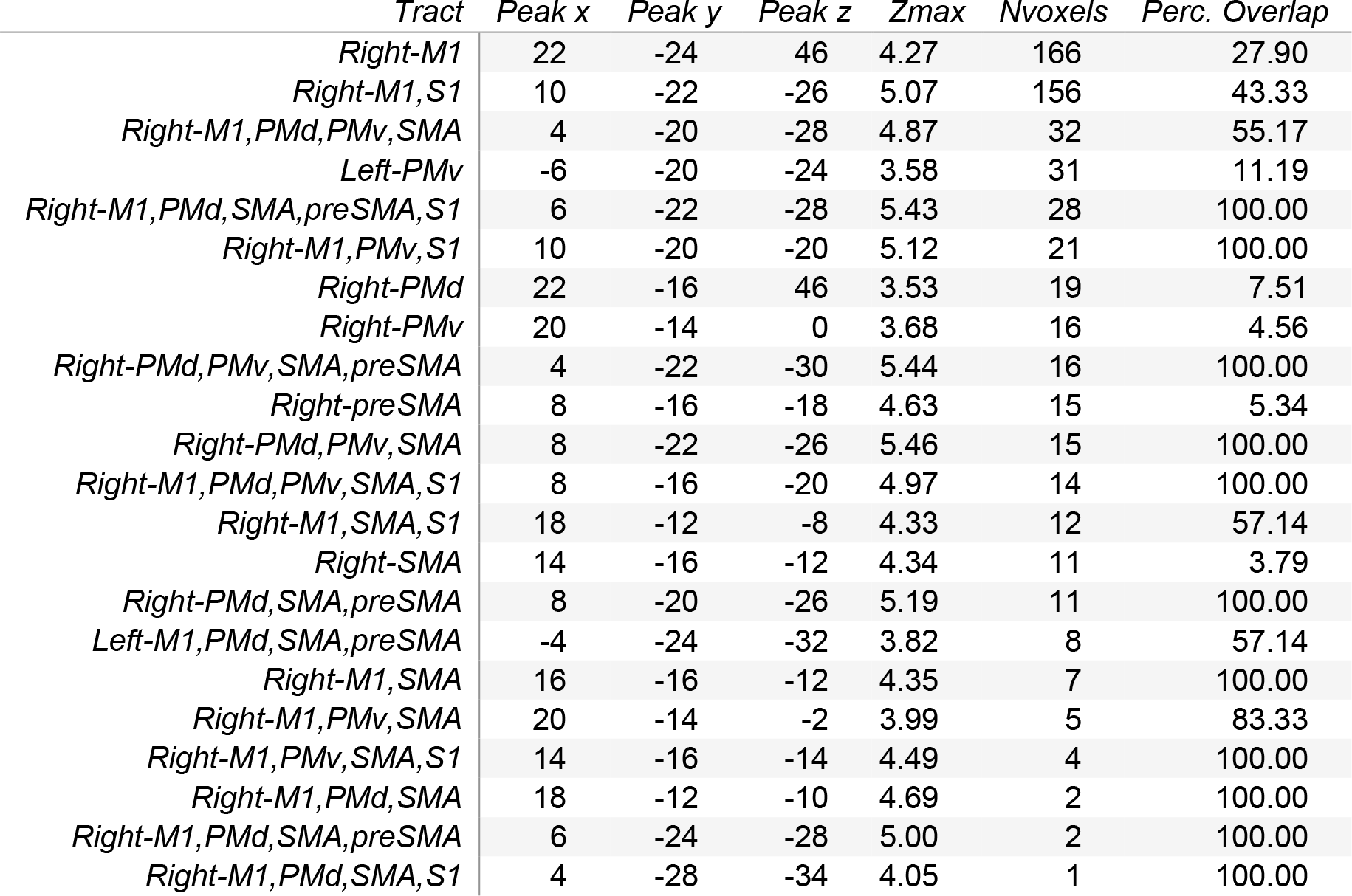
Percentage overlap between the multimodal SC brain maps and the major sensorimotor brain tracts. The labels correspond to the SMATT atlas, and only tracts attributed Z > 2 and with a percentage overlap > 10 voxels are shown. Some circuits with less than 10 voxels but with a high overlap (> 50%) are also shown.

| *Tract* | *Peak x* | *Peak y* | *Peak z* | *Zmax* | *Nvoxels* | *Perc. Overlap* |
| --- | --- | --- | --- | --- | --- | --- |
| *Right-M1* | 22 | -24 | 46 | 4.27 | 166 | 27.90 |
| *Right-M1,S1* | 10 | -22 | -26 | 5.07 | 156 | 43.33 |
| *Right-M1,PMd,PMv,SMA* | 4 | -20 | -28 | 4.87 | 32 | 55.17 |
| *Left-PMv* | -6 | -20 | -24 | 3.58 | 31 | 11.19 |
| *Right-M1,PMd,SMA,preSMA,S1* | 6 | -22 | -28 | 5.43 | 28 | 100.00 |
| *Right-M1,PMv,S1* | 10 | -20 | -20 | 5.12 | 21 | 100.00 |
| *Right-PMd* | 22 | -16 | 46 | 3.53 | 19 | 7.51 |
| *Right-PMv* | 20 | -14 | 0 | 3.68 | 16 | 4.56 |
| *Right-PMd,PMv,SMA,preSMA* | 4 | -22 | -30 | 5.44 | 16 | 100.00 |
| *Right-preSMA* | 8 | -16 | -18 | 4.63 | 15 | 5.34 |
| *Right-PMd,PMv,SMA* | 8 | -22 | -26 | 5.46 | 15 | 100.00 |
| *Right-M1,PMd,PMv,SMA,S1* | 8 | -16 | -20 | 4.97 | 14 | 100.00 |
| *Right-M1,SMA,S1* | 18 | -12 | -8 | 4.33 | 12 | 57.14 |
| *Right-SMA* | 14 | -16 | -12 | 4.34 | 11 | 3.79 |
| *Right-PMd,SMA,preSMA* | 8 | -20 | -26 | 5.19 | 11 | 100.00 |
| *Left-M1,PMd,SMA,preSMA* | -4 | -24 | -32 | 3.82 | 8 | 57.14 |
| *Right-M1,SMA* | 16 | -16 | -12 | 4.35 | 7 | 100.00 |
| *Right-M1,PMv,SMA* | 20 | -14 | -2 | 3.99 | 5 | 83.33 |
| *Right-M1,PMv,SMA,S1* | 14 | -16 | -14 | 4.49 | 4 | 100.00 |
| *Right-M1,PMd,SMA* | 18 | -12 | -10 | 4.69 | 2 | 100.00 |
| *Right-M1,PMd,SMA,preSMA* | 6 | -24 | -28 | 5.00 | 2 | 100.00 |
| *Right-M1,PMd,SMA,S1* | 4 | -28 | -34 | 4.05 | 1 | 100.00 |

**Table S5.** Anatomical description of the unimodal FC brain maps of the CCA solution that provide a maximal association with sensorimotor behavior. The labels correspond to the Desikan-Killiany atlas, and only tracts attributed Z > 2 and a percentage overlap > 50 voxels are shown.

| *Brain Area* | *Peak x* | *Peak y* | *Peak z* | *Zmax* | *Nvoxels* | *Perc. Overlap* |
| --- | --- | --- | --- | --- | --- | --- |
| *Brainstem* | 4 | -24 | -32 | 8.02 | 1287 | 35.63 |
| *ctx-rh-superiorfrontal* | 10 | 10 | 66 | 5.07 | 716 | 11.11 |
| *rh-Thalamus-proper* | 18 | -16 | 4 | 3.82 | 715 | 58.32 |
| *ctx-lh-supramarginal* | -58 | -28 | 24 | 4.08 | 616 | 19.27 |
| *lh-Thalamus-Proper* | -16 | -14 | 10 | 6.36 | 554 | 45.19 |
| *ctx-lh-superiorfrontal* | -14 | 14 | 64 | 3.45 | 329 | 5.11 |
| *ctx-lh-inferiorparietal* | -54 | -60 | 16 | 3.43 | 265 | 7.58 |
| *ctx-lh-middletemporal* | -56 | -60 | 14 | 3.44 | 261 | 10.44 |
| *ctx-lh-insula* | -28 | -18 | 10 | 3.46 | 228 | 10.06 |
| *ctx-rh-precentral* | 42 | -8 | 34 | 3.01 | 211 | 4.93 |
| *ctx-lh-precuneus* | -6 | -74 | 38 | 2.40 | 134 | 4.57 |
| *lh-Putamen* | -28 | -18 | 6 | 3.69 | 126 | 13.58 |
| *ctx-lh-bankssts* | -52 | -56 | 16 | 3.22 | 118 | 12.43 |
| *ctx-rh-postcentral* | 58 | -6 | 30 | 2.48 | 84 | 3.67 |
| *ctx-rh-supramarginal* | 54 | -26 | 26 | 2.86 | 78 | 3.07 |
| *ctx-lh-inferiortemporal* | -54 | -58 | 4 | 2.83 | 77 | 2.36 |

**Table S6.**
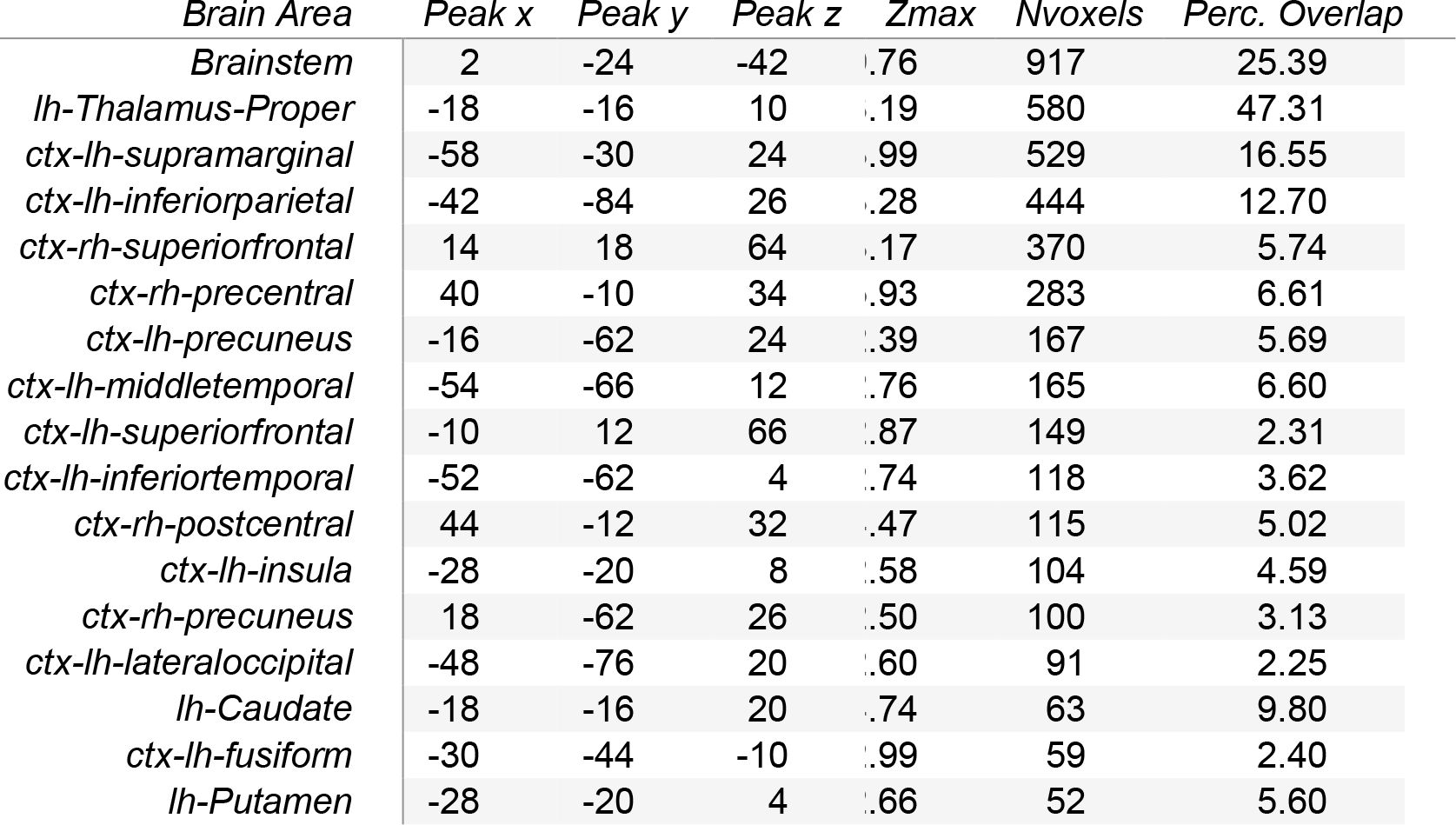
Anatomical description of the multimodal FC brain maps corresponding to the CCA solution and providing maximum association with sensorimotor behavior. Labels correspond to the Desikan-Killiany atlas, and only tracts with Z > 2 and a percentage overlap > 50 voxels are shown.

**Table S7.** Percentage overlap between the unimodal FC brain maps and the major resting state networks. The labels correspond to Yeo’s partition and only tracts with Z > 2 are shown.

| *RSN* | *Peak x* | *Peak y* | *Peak z* | *Zmax* | *Nvoxels* | *Perc. Overlap* |
| --- | --- | --- | --- | --- | --- | --- |
| *Dorsal Attention* | -18 | -14 | 10 | 6.57 | 2214 | 10.57 |
| *Limbic* | 8 | 12 | 66 | 4.80 | 990 | 2.59 |
| *Frontoparietal* | -14 | -12 | 14 | 4.82 | 867 | 3.09 |
| *Sensory-motor* | -18 | -22 | 8 | 5.24 | 824 | 3.43 |
| *DMN* | -18 | -26 | 8 | 3.72 | 535 | 3.07 |
| *Ventral Attention* | -20 | -26 | 8 | 3.85 | 278 | 1.25 |
| *Visual* | 10 | -28 | -38 | 5.96 | 138 | 0.85 |

**Table S8.** Percentage overlap between the multimodal FC brain maps and major resting state networks. Labels correspond to the Yeo’s partition and only tracts with Z > 2 are shown.

| *RSN* | *Peak x* | *Peak y* | *Peak z* | *Zmax* | *Nvoxels* | *Perc. Overlap* |
| --- | --- | --- | --- | --- | --- | --- |
| *Ventral Attention* | -18 | -16 | 12 | 8.44 | 1172 | 5.60 |
| *Sensory-motor* | -18 | -22 | 8 | 6.74 | 705 | 2.94 |
| *DMN* | -14 | -30 | 10 | 3.23 | 687 | 1.79 |
| *Dorsal Attention* | -18 | -26 | 10 | 5.75 | 652 | 3.74 |
| *Visual* | -22 | -26 | 8 | 6.01 | 557 | 2.5 |
| *Frontoparietal* | -14 | -12 | 14 | 5.57 | 389 | 1.39 |
| *Limbic* | 10 | -28 | -36 | 7.37 | 33 | 0.20 |

**Figure S1.**
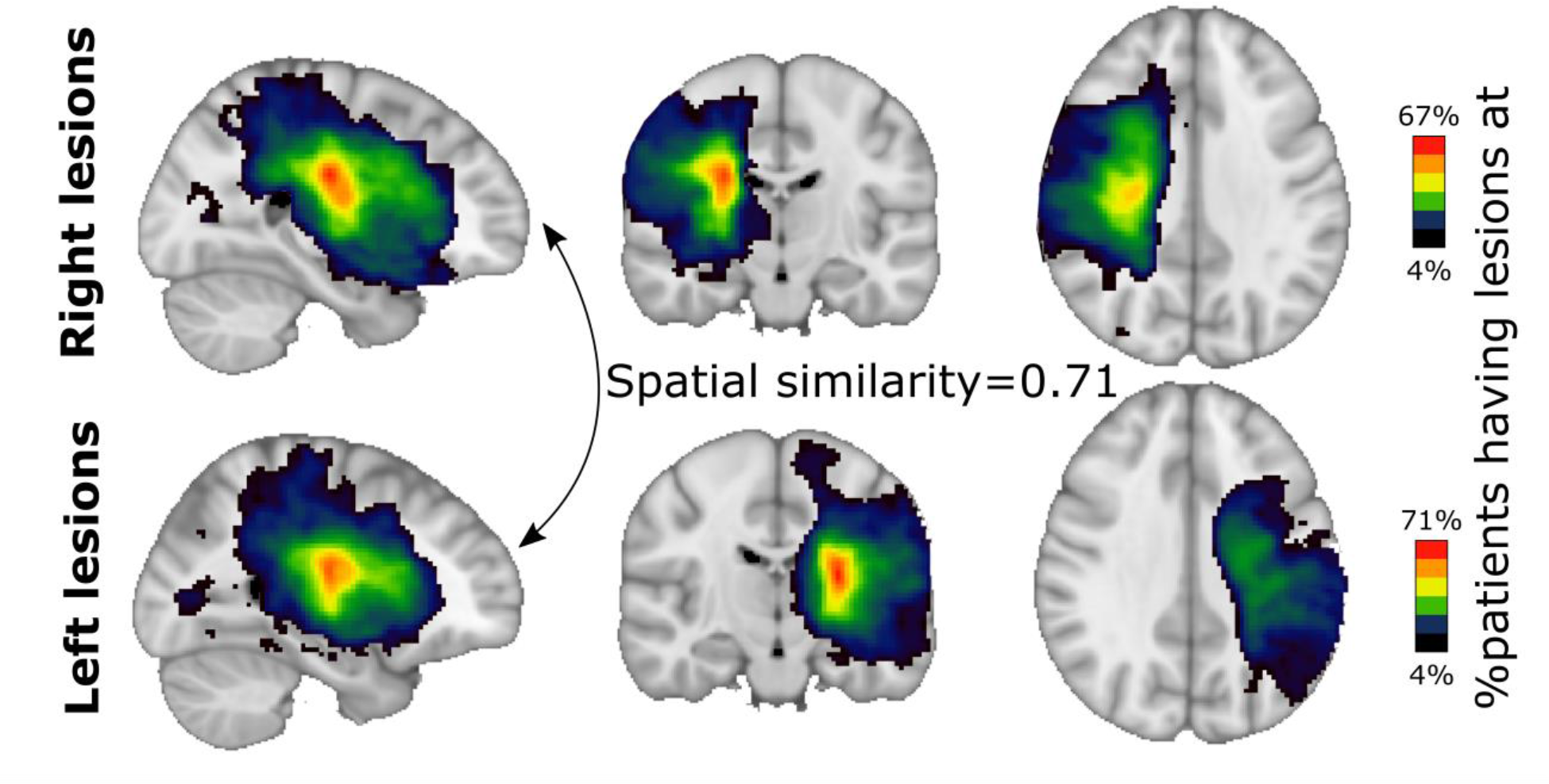
Comparison between lesion location maps of the right and left hemispheres. Top panel: The brain maps show the percentage of patients whose stroke lesion is in the right hemisphere who share the same lesion location. Lower panel: The same but for the patients whose lesion is in the left hemisphere. The spatial similarity between the two maps was 0.71, indicating a large amount of overlap between them, and indeed, the two groups of patients are difficult to differentiate when looking at the location maps (those with the lesion in the right hemisphere compared to those with the lesion in the left).

**Figure S2.**
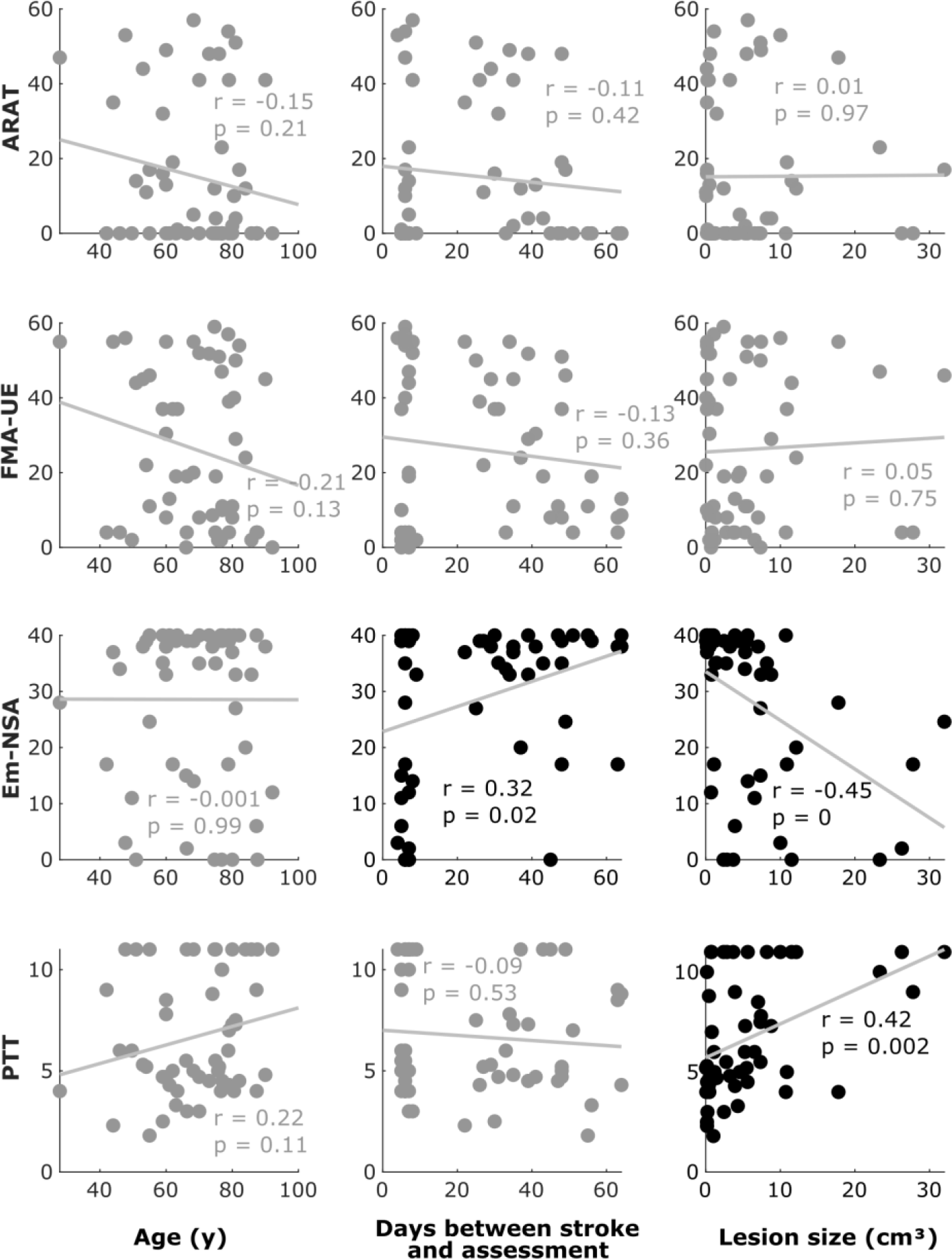
Relationship between the behavioral -sensorimotor- scores and confounders (Age, days between stroke and assessment, and lesion size). The two first rows correspond to the motor scores (ARAT, FMA-UE), and the third and fourth to the sensory scores (Em-NAS, PTT). Most of the scores show a statistical tendency to correlate with the confounders, which was regressed-out in our analyses. Scatter plots with significant correlation values (p<0.05) are highlighted in black.

**Figure S3.**
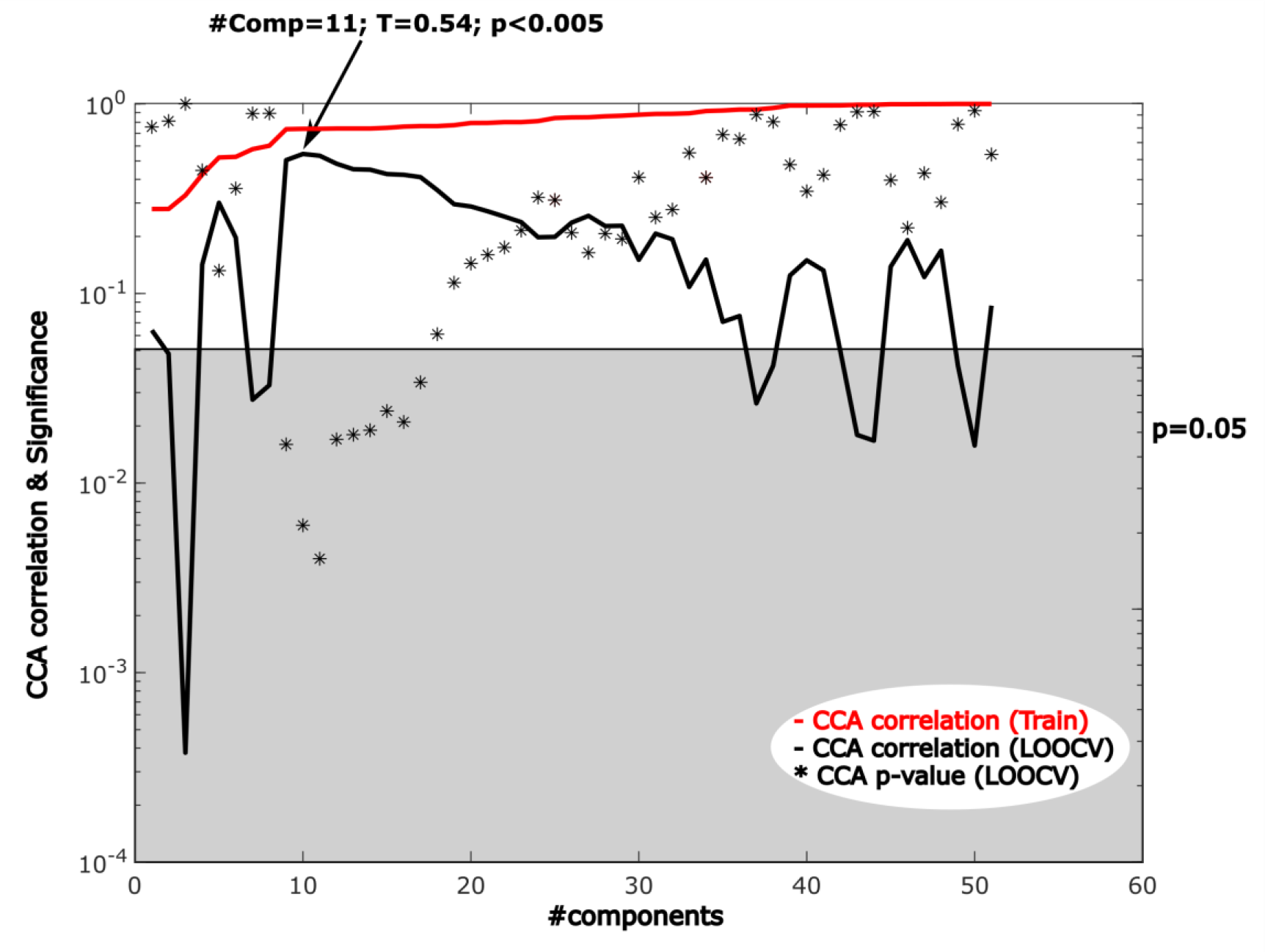
Methodological sketch for the CCA of the maximum behavioral association across imaging modalities. For each imaging modality (structural or functional) we obtained matrices of size (Npatients × Nvoxels), and we applied a PCA to obtain structural and functional components equal to the number of patients. With those components used as independent variables (input), we applied a CCA using the behavioral scores as dependent variables (output). The train curve represented here in red, shows that when the number of components increases, the input-output correlations increase monotonously, indicating over-fitting. Here, we applied leave-on-out cross-validation (LOOCV) to overcome this problem, represented here by the black curve. Maximum correlation values (represented by T) were calculated and their corresponding p-values are also reported as a measure of statistical significance. At this point of maximum correlation, the brain maps corresponding to each modality are obtained (shown in Fig. 3). Here, a log-linear representation of the data has been chosen.

**Figure S4.**
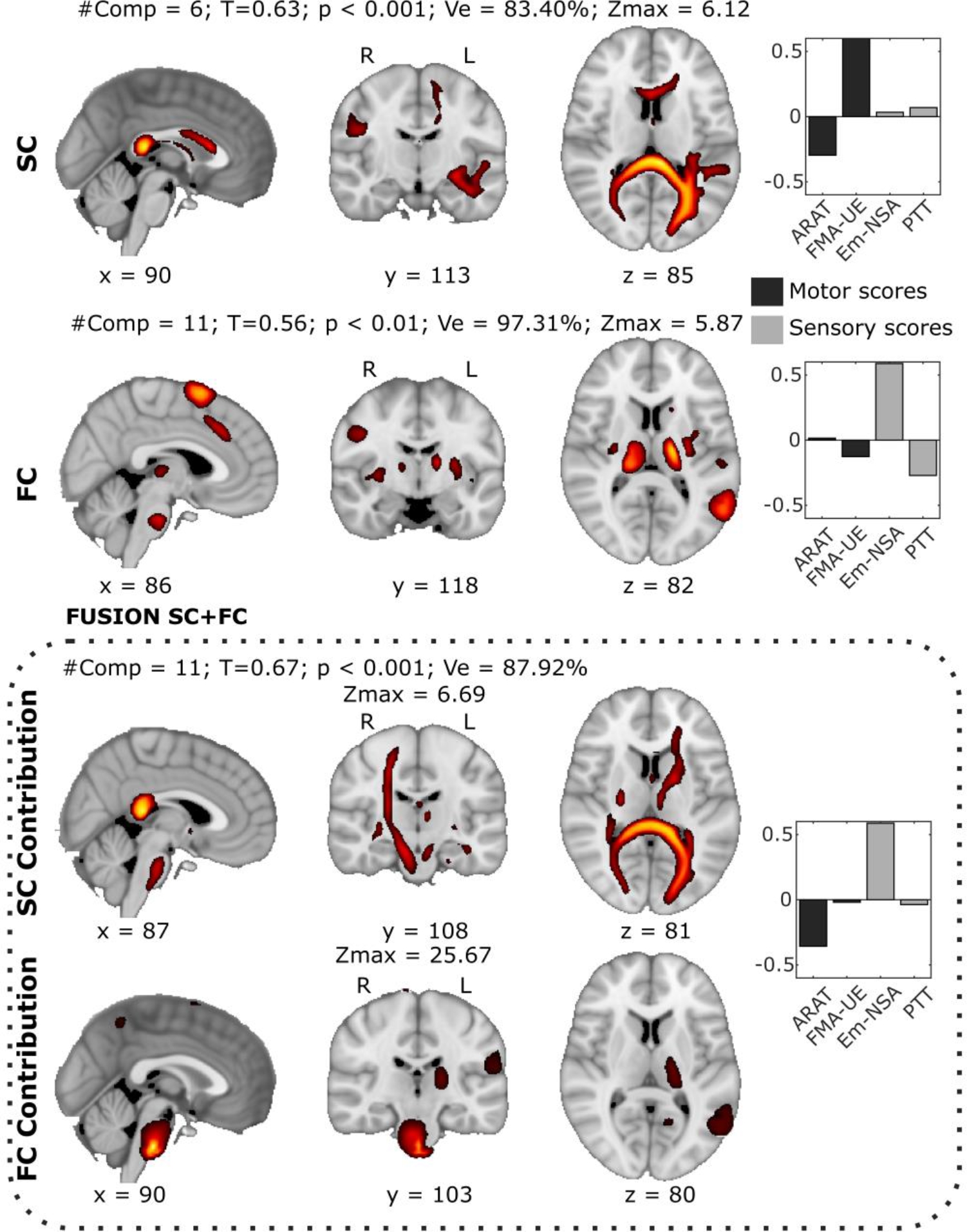
Brain maps with the maximum behavioral-association for single modality and multi- modal CCA without removing the lesion size effect. A similar analysis as in Fig. 3 but without regressing out the covariable lesion size from the behavioral scores. Major variations with respect to Fig. 3 correspond to the increased performance achieved by SC, and the different behavioral weights are represented in the right panels of each row.

**Figure S5.**
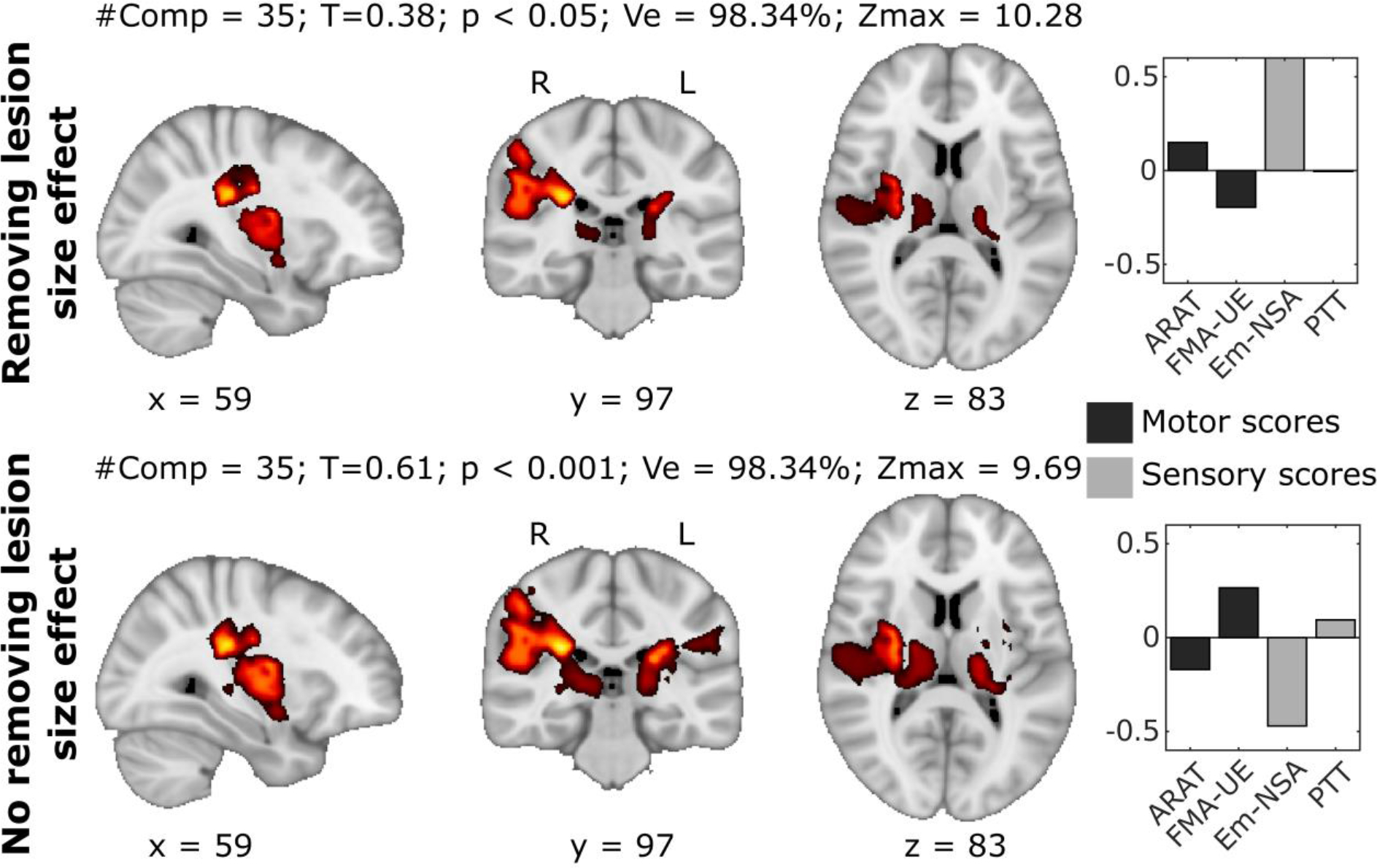
Brain maps with a maximum behavioral-association for the lesion modality with (top panel) and without (bottom panel) removing the lesion size effect. A similar analysis as in Fig. 3 but in which the PCA components have been obtained from the lesion masks and used as independent variables for the CCA analyses. Note that when the lesion size effect is not removed, the T value is considerably higher (T=0.61) than when it is removed (T=0.38).

## Footnotes

† ARAT vs. Em-NSA r = -0.03, p = 0.85; ARAT vs. PTT r = -0.08, p = 0.55; FMA-UE vs. Em-NSA r = -0.03, p = 0.81; FMA-UE vs. PTT r = -0.10, p = 0.48

‡ ARAT χ^2^ = 0.85, p = 0.36; FMA-UE χ^2^ = 0.003, p = 0.96; Em-NSA χ^2^ = 0.69, p = 0.41; PTT χ^2^ = 0.44, p = 0.51

